# Selective human tau protein expression in different clock circuits of the *Drosophila* brain disrupts different aspects of sleep and circadian rhythms

**DOI:** 10.1101/2020.12.14.422675

**Authors:** David Jaciuch, Jack Munns, Sangeeta Chawla, Seth J. Davis, Mikko Juusola

**Author notes:** Correspondence: Mikko Juusola. **Conflict of interest statement:** The authors declare no competing financial interests.

## Abstract

Circadian behavioural deficits, such as increased daytime naps and reduced night-time sleep, are common in Alzheimer’s disease and other tauopathies. But it has remained unclear whether these circadian abnormalities arise from tau pathology in either the master pacemaker or downstream neurons. Here we study this question by selectively expressing different human tau proteins in specific *Drosophila* brain circuits and monitoring locomotor activity under light-dark (LD) and in “free-running” dark-dark (DD) conditions. We show that expressing human tau proteins in the fly brain recapitulates faithfully several behavioural changes found in tauopathies. We identify discrete neuronal subpopulations within the clock network as the primary target of distinct circadian behavioural disturbances in different environmental conditions. Specifically, we show that the PDF-positive pacemaker neurons are the main site for night-activity gain and -sleep loss, whereas the non-PDF clock-neurons are the main site of reduced intrinsic behavioural rhythmicity. Bioluminescence measurements revealed that the molecular clock is intact despite the behavioural arrhythmia. Our results establish that dysfunction in both the central clock- and afferent clock-neurons jointly contribute to the circadian locomotor activity rhythm disruption in *Drosophila* expressing human tau.

**Significance Statement:** This study directly links *in vivo* human tau protein expression in region-specific *Drosophila* clock-neurons with the resulting sleep and circadian rhythm deficits to extract new knowledge of how Alzheimer’s disease and other tauopathies perturb the balance of activity and sleep. We anticipate that this novel approach will provide a useful general template for other studies of neurodegeneration in model organisms, seeking to dissect the impact of neurodegenerative disease on circadian behaviour, and further deepening our understanding of how the clock-neuron network works.

## Introduction

Alzheimer’s disease (AD) is a neurodegenerative disease that leads to progressive dementia (Salmon et al., 1999). It is characterised neuropathologically by progressive cortical neurodegeneration and the presence of extracellular amyloid plaques and intracellular tau tangles (Alzheimer, 1906; Braak and Braak, 1988; Salmon et al., 1999). The vast majority of AD patients also exhibit circadian disturbances, including increased night-time wakefulness and fragmented sleep, and reduced amplitude rhythms with phase shifts (Musiek et al., 2015).

Several animal Aβ^42^ pathology models, including 3xTG-AD mice expressing disease-linked mutant APP and tau, exhibit circadian abnormalities (Chen et al., 2014; Long et al., 2014). However, whether animal tau pathology models also display circadian dysfunction remains elusive. Circadian locomotor activity has been studied in two tau mouse models with contradictory findings (Koss et al., 2016; Stevanovic et al., 2017). Whereas, in *Drosophila*, disruption of the circadian kinase, doubletime, causes endogenous tau cleavage, resulting in circadian disturbances (Means et al., 2015).

At a cellular level, circadian rhythms emerge from interlocking ‘clock gene’ (bmal1, clock, period, cryptochrome) transcription/translation feedback loops, which produce 24-hour gene expression oscillations. Most mammalian cells exhibit circadian oscillations. However, the hypothalamic suprachiasmatic nucleus (SCN) (∼20,000 neurons) is the master pacemaker as SCN neurons receive direct photic input and entrain to light-dark (LD) cycles. They then synchronise all the other circadian oscillators through various neuronal and humoral pathways. These circadian oscillations persist in the absence of external cues (*e*.*g*. in constant darkness) accounting for “free-running” circadian activity rhythms (Buhr and Takahashi, 2013). AD patient brains show both extensive SCN cell loss (Swaab et al., 1985; Zhou et al., 1995; Swaab et al., 1998) and rhythmic but phase-shifted clocks (Cermakian et al., 2011). These findings indicate that either central clock damage, or output failure, may cause circadian dysfunction.

*Drosophila* is uniquely accessible to powerful neurogenetics and robust activity/sleep-rhythm monitoring (Fig. 1), allowing studying how tau affects the evolutionarily conserved circadian system (Bell-Pedersen et al., 2005). Its ∼150 clock-expressing neurons, organised in sLNvs, lLNvs, LPNs, LNds, DN1s, DN2s and DN3s clusters, and output neurons drive circadian locomotor activity (Fig. 1A). The neuropeptide, pigment dispersing factor (PDF), which is expressed by ∼16 lateral neurons within the sLNvs and lLNvs clusters, synchronises all the clock-neurons. Specifically, the sLNVs are the master pacemakers, as they control the speed of “free-running” behavioural and molecular rhythms in a PDF-dependent manner (Dubowy and Sehgal, 2017).

**Figure 1.**
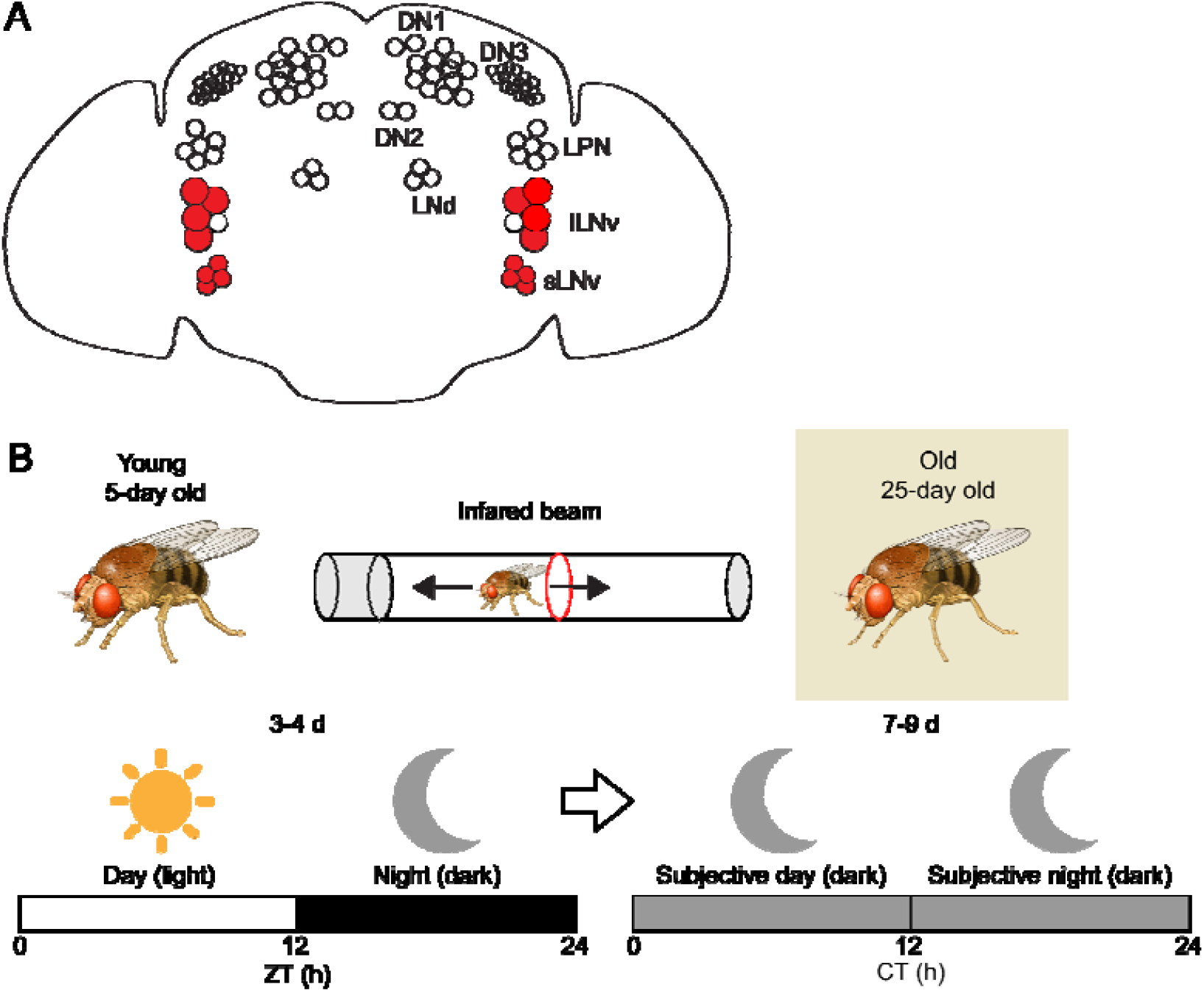
(A) **Clock system in the Drosophila brain.** In the fly brain, ∼150 neurons express a molecular clock. The ∼150 clock-neurons form clusters (sLNv, lLNv, LNd, LPN, DN1, DN2 and DN3). Different neurons serve different functions and respond to different environmental conditions. The PDF-positive clock-neurons (shown in red) are the master pacemakers as they synchronise all the clock-neurons. (B) **Experimental protocol**. Individual male 5-(young) and 25-day old (old) flies were placed in glass tubes in a Drosophila Activity Monitor (in a light- and temperature-controlled incubator) which counts the number of times a fly breaks a beam bisecting the tube. Locomotor activity was first recorded in a 12 h light:dark (LD) cycle for three-four days followed by seven-nine days of continuous darkness (DD).

*Drosophila* tauopathy models, in which human wild-type or frontotemporal dementia (FTD)-linked mutant tau is ectopically expressed in the developing fly brain or visual system reproduce many of the behavioural and neurophysiological changes seen in human AD patients, including adult-onset progressive neurodegeneration, a reduced lifespan and learning and memory deficits (Wittmann et al., 2001; Jackson et al., 2002; Mershin et al., 2004; Nishimura et al., 2004). However, hitherto no studies have compared the circadian and sleep disturbances in *Drosophila* expressing different human tau proteins in discrete brain circuits at various ages.

In this study, we discovered both isoform- and region-specific differences in tau-induced circadian behavioural abnormalities in different light conditions. We identified the PDF expressing-neurons as the main site of activity gain and sleep loss, affecting the LD conditions’ night component. The non-PDF clock-neurons were found to be the main site of reduced intrinsic behavioural rhythmicity. Through bioluminescence measurements, we showed that the molecular clock is functional in tau-expressing flies despite the behavioural arrhythmia. These results suggest that the circadian and sleep phenotypes in human tau-expressing *Drosophila* emerge from disrupted communication between the central clock and downstream clock-neurons and the clock-neurons output neurons, rather than from damage to the master pacemaker. As the tau-expressing flies’ behavioural changes mirror those seen in human AD patients, we suggest that tau-mediated clock neuronal dysfunction drives the sleep and circadian phenotypes in both flies and humans. We further suggest that both isoform- and region-specific effects contribute to the discrete circadian and sleep phenotypes in distinct tauopathies.

## Materials and Methods

### Fly stocks

Flies were raised on standard cornmeal food under a 12 h light:12 h dark (LD) cycle at 25 °C and 70 % humidity. All lines were backcrossed at least five generations to the Canton-S wild-type stock. The following lines were used in the study: Canton-S (#1), Elav^c155^-Gal4 (#458) and UAS-human 2n4r tau^WT^1.13 (#51362). These were obtained from the Bloomington *Drosophila* Stock Centre. UAS-human 0n4r tau^R406W^ (Wittmann et al., 2001) was a gift from Dr Mel Feany (Harvard Medical School, USA). Tim-Gal4, Pdf-Gal4 (Kaneko and Hall, 2000) and BG-luc (Stanewsky et al., 1997) flies were kindly provided by Dr Ralf Stanewsky (University of Münster, Germany). Pdf-Gal80 (Stoleru et al., 2004) was a gift from Dr Charlotte Forster (University of Würzburg, Germany).

### Locomotor behaviour assay

Adult males were collected within a few hours of eclosion and ≤20 were aged on standard food and tipped onto new food every two-three days. Individual male flies were placed in 65 x 5 mm glass tubes, containing a small amount of 5 % sucrose and 2 % agarose dissolved in water. Locomotor activity was recorded with *Drosophila* Activity Monitors (DAMs; TriKinetics, USA), which count the number of times the fly breaks an infrared beam bisecting the tube (Fig. 1B). Monitors were placed in a light- and temperature-controlled (25°C) incubator (Panasonic Mir c155, Japan). By placing a small beaker of water inside the incubator, humidity was kept between 50-70 %.

Locomotor activity of both young (5-day old) and old (25-day old) flies was measured for three-four days in 12 h LD cycles followed by seven-nine days in constant darkness (DD). Locomotor activity and sleep profiles were produced from three-four consecutive LD days or seven-nine consecutive DD days data. Daytime and night-time activity is the total number of beam breaks during the 12 h light or dark period, respectively, averaged across at least three consecutive days. Sleep was defined as a period of at least five minutes of inactivity (Hendricks et al., 2000; Shaw et al., 2000). Sleep analysis was conducted using a custom-written Excel macro (Donlea et al., 2014). Daytime and night-time sleep is the total sleep during the 12 h light or dark period, respectively, averaged across at least three consecutive days.

The DD activity data were analysed by Lomb-Scargle periodogram analysis (van Dongen and Chapman, 1999) with the Actogram J program (Schmid et al., 2001; available at http://imagej.net/ActogramJ) to determine the rhythmicity and period (*i*.*e*. the length of the intrinsic day) of behavioural rhythms. The rhythmicity (power) was defined as the amplitude of the peak only for flies deemed rhythmic. Flies were determined to be rhythmic or arrhythmic based upon the presence or absence of a peak as convention above the p<0.05 significance level. Only rhythmic flies were used to calculate the behavioural period. For more detailed information, see (Kauranen et al., 2012).

### Luciferase Assay

Both young (5-day-old) and old (25-day-old) flies were placed individually in every other well of a 96-well white microtiter plate (Perkin Elmer, USA) in which each well contained 200 µl of a 5 % sucrose, 1 % agarose and 15mM D-luciferin (SynChem, USA) solution. Plates were first exposed to a 12 h LD cycle for three days at 25 °C. Plates were then loaded into a TopCount Scintillation Counter (Packard, USA) and bioluminescence was measured for four days in continuous darkness (Stanewsky et al., 1997). The TopCount Scintillation Counter was housed in a 25 °C room and was modified as described (Anwer et al., 2014). Both relative amplitude error (RAE) and period were calculated using BRASS (Locke et al., 2005). RAE is a measure of rhythm robustness that ranges from 0 (a perfect fit to the wave) to 1 (no fit). As a convention, flies with ∼0.7 ≤ RAE ≤ 1 were classed as rhythmic.

### Statistics

Samples sizes are reported in figures. Box plots show median with interquartile range and the 10 and 90 percentiles as whiskers. Flies which did not survive the experiment were excluded from the analysis. All datasets were tested for normal or lognormal distribution by a Kolmogorov-Smirnov test. Activity and power datasets were lognormally distributed. Therefore log-transformed data were analysed by 1-way ANOVA (single factor, genotype) or 2-way ANOVA (two factors, genotype and age), followed by post-hoc tests. Multiple comparisons after ANOVA were performed by a Tukey HSD test.

Sleep and period datasets were neither normally or lognormally distributed. Therefore, we chose to use non-parametric tests, rather than parametric tests (t-test and ANOVA) that assume normal distribution. Sleep and period datasets were analysed by Kruskal Wallis ANOVA (single factor, genotype). Sleep and period datasets with two factors were analysed in two ways. First, Kruskal-Wallis ANOVA followed by post-hoc tests were used to check for differences between different genotypes of the same age. Multiple comparisons after ANOVA were performed by a Dunn’s test. Second, a Mann-Whitney U-test was used to check for differences between different ages of the same genotype.

Bioluminescence datasets were analysed by Mann-Whitney U-tests to check for differences between different genotypes of the same age or different ages of the same genotype, as they did not follow a normal distribution.

P levels are indicated as non-significant (ns) p > 0.05, * p < 0.05, ** p < 0.001 or *** p < 0.0001.

## Results

The R406W tau mutation found in frontotemporal dementia and parkinsonism linked to chromosome 17 (FTDP17) causes a hereditary tauopathy clinically resembling AD, associated with early-onset and rapid progression (Hutton et al., 1998; Saito et al., 2002). As different tau proteins are associated with distinct tauopathies with specific clinical symptoms (Josephs, 2017), they may precipitate discrete behavioural changes when studied in isolation.

### Pan-neuronal tau expression disturbs activity and sleep under LD conditions

To investigate how tau affects circadian behaviour, full-length human wild-type (WT) (2n4r isoform) (Jackson et al., 2002) and mutant (R406W) (0n4r isoform) (Wittmann et al., 2001) tau were expressed in *Drosophila* pan-neurally, using the Gal4/UAS system (Elav^c155^-Gal4 driver) (Brand and Perrimon, 1993). We then recorded locomotor activity in young (5-day-old) and old (25-day-old) tau-expressing flies under a 12 h LD cycle and in continuous darkness (Fig. 1B). Examining both young and old flies enabled us to assess whether behavioural changes were affected by ageing or progressive. As a high mortality rate in the Elav>tau flies beyond 40-days of age confounded our observations of circadian behaviour, we did not monitor activity rhythms in older individuals.

Under LD conditions, both the Gal4- and UAS-control flies exhibited wild-type circadian behaviour (Fig. 2A, left). The activity profiles contained morning and evening activity peaks, centred around the light transitions (lights on: zeitgeber time (ZT) = 0; lights off: ZT = 12), separated by a midday siesta and a period of consolidated sleep during the night (Dubowy and Sehgal, 2017). The Elav>2n4r tau^WT^ and Elav>0n4r tau^R406W^ flies, on the other hand, showed normal bimodal activity profiles, but elevated baseline activity, particularly during the second half of the night (Fig. 2A, middle and right). Overall, daytime activity levels (Fig. 2B) were indistinguishable in the Elav>tau and control flies, except for a small statistically significant reduction in the old Elav>0n4r tau^R406W^ flies. However, in clear contrast, the night activity levels (Fig. 2C) of both the young and old Elav>tau flies were greatly increased, with both tau proteins producing a similar gain in night activity. Between the young and old age groups, we found no statistically significant age-related differences in the daytime (Fig. 2B) and night-time (Fig. 2C) activity in either the Elav>tau or control flies.

**Figure 2.**
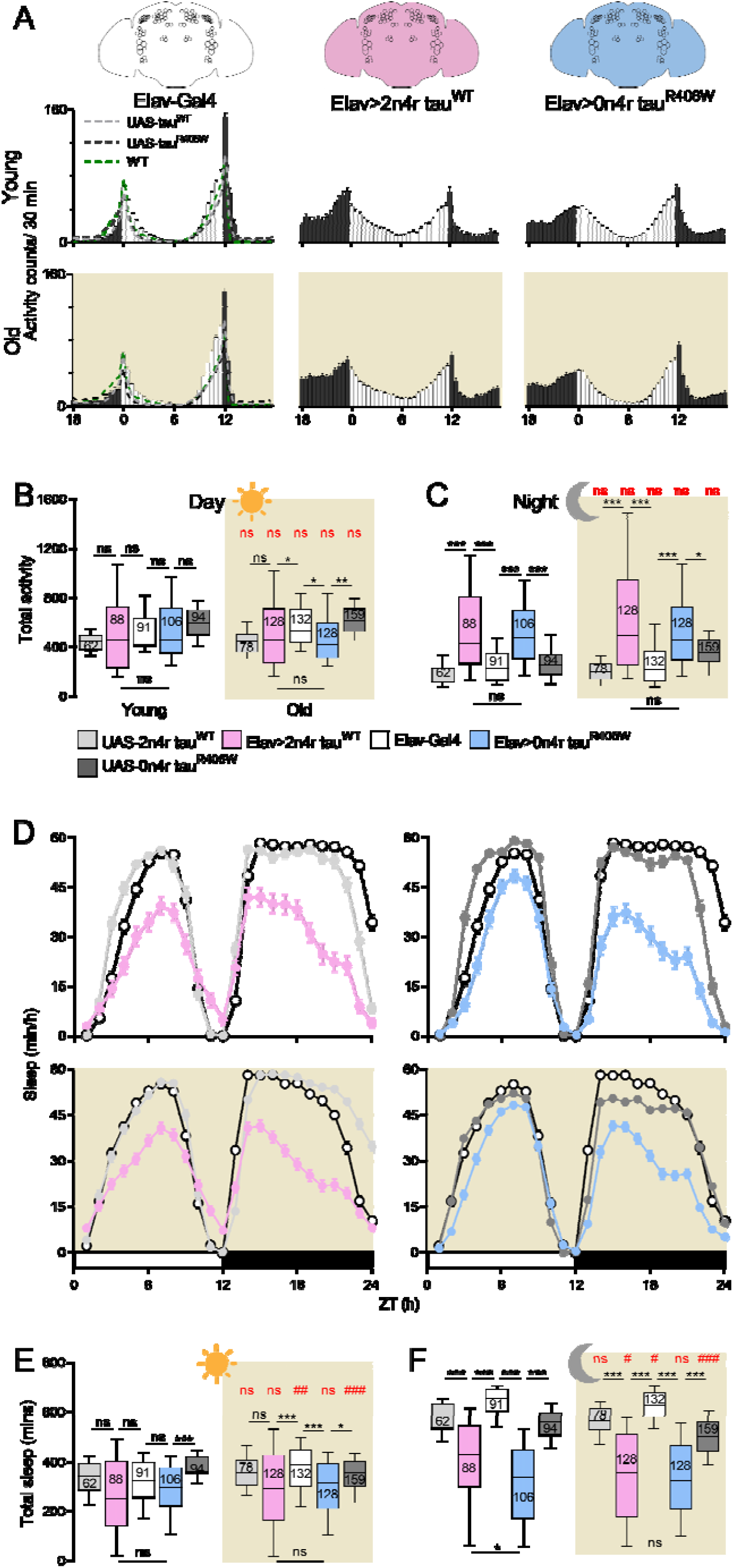
Pan-neural tau-expression alters activity and sleep levels under a 12 h light: 12h dark (LD) cycle. **(A)** Activity histograms of 5-(young) (top) and 25-day old (old) (bottom, brown shading) control, Elav>2n4r tau^WT^ and Elav>0n4r tau^R406W^ flies. All genotypes show normal bimodal activity rhythms. Elav>tau flies’ basal activity level is elevated. Bars/lines show mean ± SEM in 30 min bins (lights-on = white bars, lights-off = dark-grey bars). (**B-C**) Quantifying daytime and night-time activity, respectively. Elav>tau expression greatly increases night-time activity relative to controls. Multiple comparisons between different genotypes of the same age (black asterisks) and different ages of the same genotype (red number symbols) by 2-way ANOVA and post-hoc Tukey HSD tests with log-transformed data. (**D**) Sleep profiles of young (top) and old (bottom, brown shading) Elav>2n4r tau^WT^ and Elav>0n4r tau^R406W^ flies compared to relevant controls. All genotypes show a normal bimodal profile. Symbols show mean ± SEM. (**E-F**) Quantifying daytime and night-time sleep, respectively. Elav>tau expression greatly reduces daytime and night-time sleep relative to controls. Multiple comparisons between different genotypes of the same age (black asterisks) by Kruskal-Wallis ANOVA and post-hoc Dunn’s tests. Comparisons between different ages of the same genotype (red number symbols) by Mann-Whitney U-tests. Materials and Methods describe statistics. Box plots show median with 2^nd^ and 3^rd^ quartiles and 10 and 90 percentiles as whiskers. n = 62-159 flies from 4-8 independent experiments.

Next, we investigated whether the elevated night activity in the Elav>tau flies coincides with sleep loss by examining their sleep. The sleep profiles revealed that the Elav>tau flies seem to sleep less throughout the day and night (Fig. 2D). However, the daytime sleep loss fell just short of significance in all except the old Elav>0n4r tau^R406W^ flies, which only just reached the significance threshold (Fig. 2E). In contrast, the night-time sleep loss was highly significant with respect to the controls in both young and old flies (Fig. 2F). These results collectedly showed that broad neuronal human tau expression promotes activity during the night and suppresses sleep throughout the day and night.

### Pan-neural tau expression disrupts “free-running” circadian behavioural rhythms

Human patients with AD often have circadian rhythm defects, which result in disrupted body temperature and activity rhythms (Satlin et al., 1995; Harper et al., 2001). Therefore, to assess internal clock function, we next monitored the locomotor activity of the Elav>tau and control flies in the absence of external cues, in continuous darkness. In such conditions, the Gal4- and UAS-control flies maintained wild-type daytime activity and night-time inactivity patterns with a period of nearly 24 h (Fig. 3A, left). Both the Elav>2n4r tau^WT^ (Fig. 3A, middle) and Elav>0n4r tau^R406W^ (right) flies were similarly more day-than night-active, but the distinction between day activity and night inactivity was less obvious, as relative night activity seemed to be elevated.

**Figure 3.**
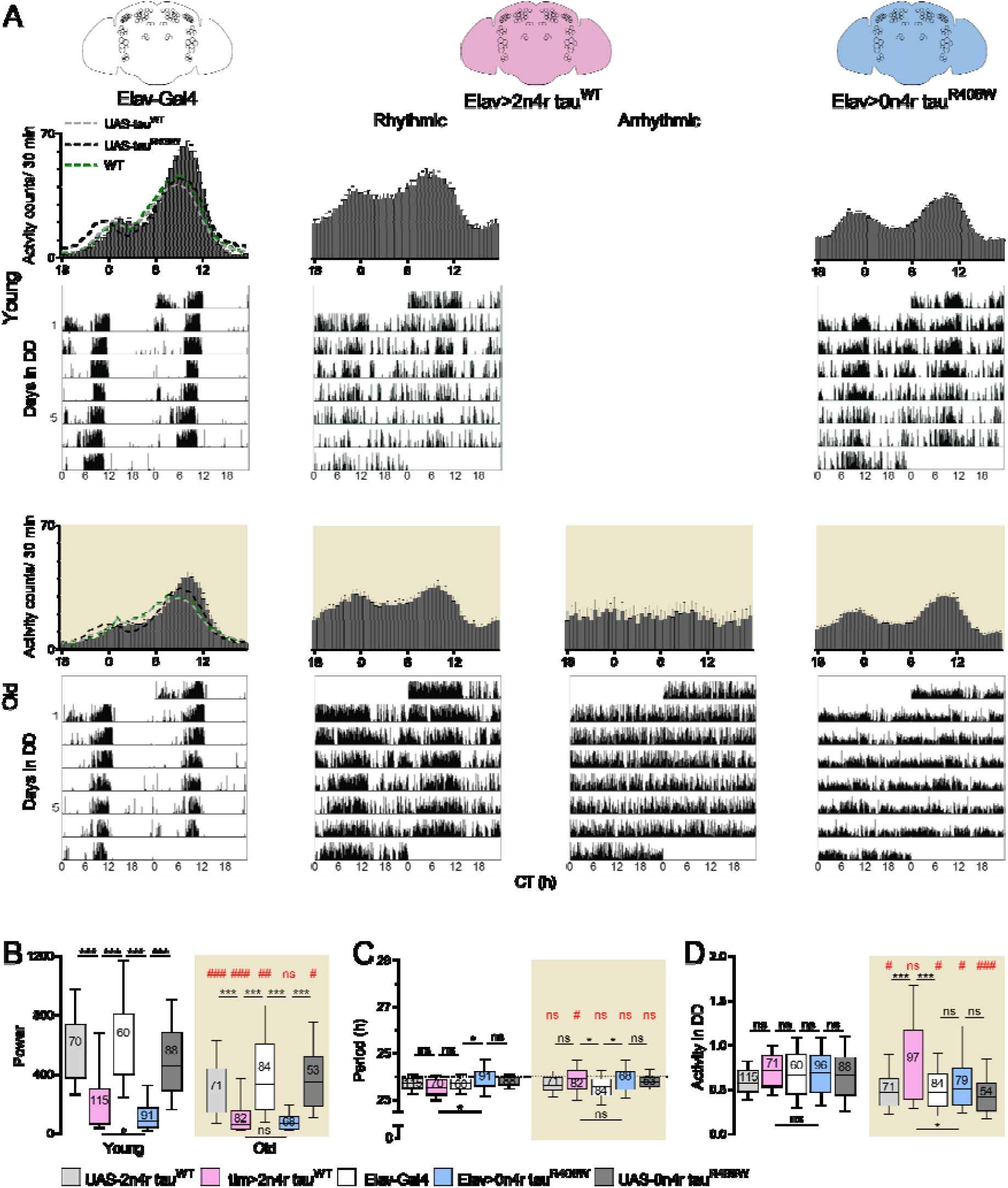
Pan-neural tau expression produces arrhythmia in “free-running” DD conditions. (**A**) Activity histograms and representative double-plotted actograms for 5-(young) (top) and 25-day old (old) (bottom, brown shading) control, Elav>2n4r tau^W^^T^ and Elav>0n4r tau^R406W^ flies. Control flies show robust daytime activity and night-time inactivity rhythms, with a ∼24 h period (left). Elav>tau flies show either weak rhythms with a normal period (middle and right) or are arrhythmic (middle). Table 1 gives the percentage of rhythmic flies. Bars/lines show mean ± SEM in 30 min bins. (**B**) Pan-neuronal tau expression greatly reduces DD behavioural rhythmicity, determined by Lomb-Scarle analysis. (**C**) Elav>tau expression does not affect the behavioural period in DD. (**D**) Elav>2n4r tau^WT^ expression produces age-related DD hyperactivity. But Elav>0n4r tau^R406W^ expression does not affect overall DD activity. (**B, D**) Multiple comparisons between different genotypes of the same age (black asterisks) and different ages of the same genotype (red number symbols) by 2-way ANOVA and post-hoc Tukey HSD tests with log-transformed data. (**C**) Multiple comparisons between different genotypes of the same age (black asterisks) by Kruskal-Wallis ANOVA and post-hoc Dunn’s tests. Comparisons between different age of the same genotype (red number symbols) by Mann-Whitney U-tests (for further details see Materials and Methods). n = 54-115 flies from 4-6 independent experiments.

The strength of the circadian rhythms was assessed by Lomb-Scargle periodogram analysis (van Dongen and Chapman, 1999). We found that DD locomotor behaviour’s rhythmicity was greatly reduced in both the Elav>2n4r tau^WT^ and Elav>0n4r tau^R406W^ flies compared to the age-matched controls. In the young flies, we observed a significantly larger reduction in behavioural rhythmicity in the Elav>0n4r tau^R406W^ flies relative to the Elav>2n4r tau^WT^ flies. However, we found a statistically significant age-related decline in DD rhythmicity in both the Elav>2n4r tau^WT^ and control flies, but not in the Elav>0n4r tau^R406W^ flies. Consequently, in the old flies, Elav>2n4r tau^WT^ and Elav>0n4r tau^R406W^ expression produced a similar reduction in circadian rhythmicity (Fig. 3B). A subpopulation of the old Elav>tau flies developed arrhythmia, being active around the clock (≤10 %). But in comparison, 100 % of the control flies remained rhythmic (Table. 1). An average activity histogram and representative double-plotted actogram for a rhythmic and arrhythmic Elav>2n4r tau^WT^ fly is shown in Fig. 3A (middle). The pan-neuronal expression of tau had no effect on the behavioural period in DD at both ages analysed (Fig. 3C). Together, these results indicated that broad neuronal tau expression reduces circadian rhythmicity without altering the behavioural period in DD.

**Table 1.** Circadian behaviour of flies in DD. Power and period were determined by Lomb-Scargle periodogram analysis. Flies were defined as rhythmic based upon the presence of a peak above the 0.05 significance line. The rhythmic percentage is the number of rhythmic flies/ numbers of tested flies as a percentage. Survival percentage is the number of flies that survived to the end of the experiment/ number of flies that started the experiment as a percentage.

| Genotype | Age (d) | n | Power |  | Period (h) |  |  | Rhythmic % | Activity in DD |  | Survival % | Colour |
| --- | --- | --- | --- | --- | --- | --- | --- | --- | --- | --- | --- | --- |
| | | | median | mean ± SEM | median | mean ± SEM | range | | median | mean ± SEM | | |
| Elav-Gal4 | 5 | 60 | 574 | 634 ± 43 | 23.7 | 23.7 ± 0.04 | 1.8 | 100 | 0.66 | 0.69 ± 0.04 | 94 | |
| Elav-Gal4 | 25 | 84 | 330 | 410 ± 34 | 23.6 | 23.6 ± 0.10 | 4.7 | 100 | 0.46 | 0.51 ± 0.03 | 93 | |
| UAS-2n4r tau<sup>WT</sup> | 5 | 11<br>5 | 529 | 597 ± 31 | 23.7 | 23.7 ± 0.03 | 1.5 | 100 | 0.57 | 0.60 ± 0.02 | 96 | |
| UAS-2n4r tau<sup>WT</sup> | 25 | 71 | 226 | 293 ± 28 | 23.6 | 23.6 ± 0.06 | 3.3 | 100 | 0.46 | 0.49 ± 0.03 | 92 | |
| Elav-Gal4/UAS-2n4r tau<sup>WT</sup> | 5 | 71 | 156 | 237 ± 29 | 23.5 | 23.5 ± 0.05 | 2.2 | 98 | 0.71 | 0.74 ± 0.03 | 89 | |
| Elav-Gal4/UAS-2n4r tau<sup>WT</sup> | 25 | 97 | 58 | 118 ± 15 | 23.9 | 24.0 ± 0.15 | 11.9 | 84 | 0.61 | 0.83 ± 0.06 | 88 | |
| UAS-0n4r tau<sup>R406W</sup> | 5 | 88 | 454 | 527 ± 34 | 23.6 | 23.8 ± 0.02 | 1.0 | 99 | 0.66 | 0.68 ± 0.04 | 95 | |
| UAS-0n4r tau<sup>R406W</sup> | 25 | 54 | 346 | 382 ± 35 | 23.5 | 23.8 ± 0.05 | 1.5 | 98 | 0.41 | 0.46 ± 0.04 | 89 | |
| Elav-Gal4/UAS-0n4r tau<sup>R406W</sup> | 5 | 96 | 81 | 142 ± 18 | 23.6 | 23.9 ± 0.09 | 8.2 | 95 | 0.69 | 0.71 ± 0.03 | 86 | |
| Elav-Gal4/UAS-0n4r tau<sup>R406W</sup> | 25 | 79 | 65 | 81 ± 8 | 23.4 | 24.0 ± 0.12 | 6.4 | 86 | 0.50 | 0.59 ± 0.04 | 59 | |
| tim-Gal4 | 5 | 92 | 335 | 447 ± 33 | 23.8 | 23.8 ± 0.03 | 1.4 | 100 | 0.59 | 0.66 ± 0.03 | 96 | |
| tim-Gal4 | 25 | 67 | 275 | 317 ± 32 | 23.6 | 23.6 ± 0.06 | 2.9 | 100 | 0.53 | 0.59 ± 0.02 | 87 | |
| tim-Gal4/2n4r tau<sup>WT</sup> | 5 | 46 | 60 | 74 ± 7 | 24.1 | 24.7 ± 0.25 | 8.4 | 95 | 1.15 | 1.10 ± 0.06 | 100 | |
| tim-Gal4/2n4r tau<sup>WT</sup> | 25 | 77 | 24 | 30 ± 2 | 24.7 | 25.1 ± 0.35 | 12.4 | 70 | 0.57 | 0.61 ± 0.04 | 96 | |
| tim-Gal4/UAS-0n4r tau<sup>R406W</sup> | 5 | 88 | 68 | 86 ± 7 | 24.2 | 24.2 ± 0.13 | 11.7 | 98 | 0.81 | 0.82 ± 0.03 | 97 | |
| tim-Gal4/UAS-0n4r tau<sup>R406W</sup> | 25 | 50 | 37 | 50 ± 5 | 24.7 | 24.9 ± 0.29 | 12.2 | 98 | 0.84 | 0.93 ± 0.05 | 98 | |
| Pdf-Gal4 | 5 | 71 | 508 | 560 ± 25 | 23.8 | 23.8 ± 0.02 | 2.0 | 100 | 0.55 | 0.58 ± 0.02 | 96 | |
| Pdf-Gal4 | 25 | 60 | 228 | 303 ± 33 | 23.8 | 23.8 ± 0.06 | 2.6 | 100 | 0.42 | 0.50 ± 0.04 | 89 | |
| Pdf-Gal4/2n4r tau<sup>WT</sup> | 5 | 55 | 519 | 533 ± 29 | 24.5 | 24.5 ± 0.05 | 1.6 | 100 | 1.00 | 1.07 ± 0.05 | 98 | |
| Pdf-Gal4/2n4r tau<sup>WT</sup> | 25 | 31 | 199 | 230 ± 25 | 24.5 | 24.5 ± 0.06 | 1.5 | 100 | 0.90 | 0.97 ± 0.07 | 97 | |
| Pdf-Gal4/UAS-0n4r tau<sup>R406W</sup> | 5 | 10<br>9 | 189 | 233 ± 16 | 24.3 | 24.3 ± 0.03 | 1.8 | 100 | 0.83 | 0.85 ± 0.02 | 97 | |
| Pdf-Gal4/ UAS-0n4r tau<sup>R406W</sup> | 25 | 27 | 183 | 183 ± 20 | 24.3 | 24.3 ± 0.06 | 1.2 | 100 | 0.99 | 1.03 ± 0.05 | 84 | |
| Elav-Gal4, Pdf-Gal80 | 5 | 48 | 556 | 604 ± 63 | 23.4 | 23.4 ± 0.04 | 1.3 | 100 | 0.90 | 0.94 ± 0.07 | 100 | |
| Elav-Gal4, Pdf-Gal80/UAS-0n4r tau<sup>R406W</sup> | 5 | 32 | 67 | 105 ± 21 | 23.7 | 23.8 ± 0.14 | 4.9 | 100 | 0.71 | 0.85 ± 0.08 | 91 | |

Because ubiquitous neuronal human tau expression in *Drosophila* can cause motor deficits (Ali et al., 2012), it was possible that the DD arrhythmic phenotype was related to reduced activity levels. However, we found no statistically significant differences in overall DD activity between the Elav>0n4r tau^R406W^ and control flies, at both ages analysed. There was a small statistically significant decline in DD activity between the young and old age groups in both the Elav>0n4r tau^R406W^ and control flies. Intriguingly, Elav>2n4r tau^WT^ expression resulted in age-related DD hyperactivity; activity levels were normal in the young, but greatly increased in the old, compared to age-matched controls (Fig. 3D). Therefore, the Elav>tau flies’ behavioural arrhythmia was not an artefact of reduced activity levels.

### Tau expression specifically in the clock network alters activity and sleep under LD conditions

Post-mortem human AD patient brains show extensive SCN cell loss, suggesting central clock damage might account for the circadian behavioural deficits (van Dongen and Chapman, 1999). Therefore, we next assessed the consequences of restricting tau expression to the fly clock network. To achieve this, we used tim-Gal4 and Pdf-Gal4 to drive tau expression in all clock cells (∼150 neurons) or exclusively in the PDF-positive pacemaker neurons (∼16 neurons), respectively, and recorded locomotor activity under both LD and DD conditions.

First, we examined the effects of pan-clock tau expression on locomotor behaviour under LD conditions. Both the tim>2n4r tau^WT^ (Fig. 4Aiii) and tim>0n4r tau^R406W^ (iv) flies showed normal bimodal activity rhythms with elevated basal activity, particularly during the second half of the night, compared to the Gal4-and UAS-control flies (i-ii). For both young and old, we found significantly increased total night activity in the tim>tau flies (Fig. 4C). Tim>tau expression also yielded flies, which exhibited dramatically reduced night sleep in both age groups (Figs. 4Di-ii, F). In the young flies, tim>2n4r tau^WT^ and tim>0n4r tau^R406W^ expression produced a similar activity gain and sleep loss at night. However, the phenotype was not stable in the tim>2n4r tau^WT^ flies as they aged. Therefore, in the old flies, the night-activity gain and -sleep loss was significantly smaller in the tim>2n4r tau^WT^ flies relative to the tim>0n4r tau^R406W^ flies (Figs. 4C, F).

**Figure 4.**
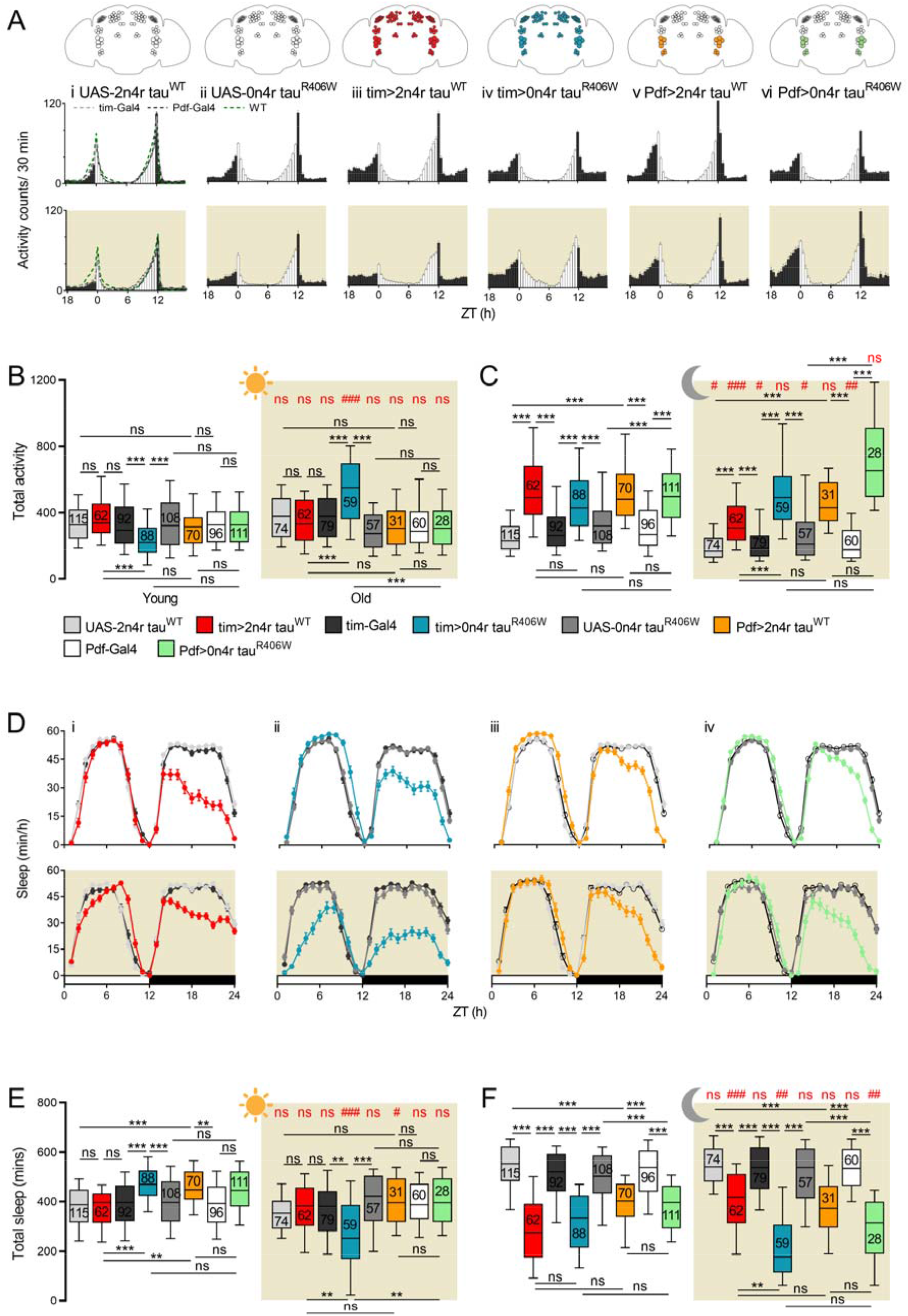
Region- and isoform-specific differences of clock-specific tau expression on activity and sleep levels under LD conditions. (**A**) Activity histograms of 5-(young) (top) and 25-day old (old) (bottom, brown shading) (i-ii) UAS control, (iii) tim>2n4r tau^WT^, (iv) tim>0n4r tau^R406W^, (v) Pdf>2n4r tau^WT^ and (vi) Pdf>0n4r tau^R406W^ flies. All genotypes show a normal bimodal activity profile. Elevated baseline activity in both the tim>tau and Pdf>tau flies. (**B-C**) Quantifying daytime and night-time activity, respectively. Tim>tau and Pdf>tau expression does not affect daytime activity (except for tim>0n4r tau^R406W^ expression), but greatly increases night-activity, relative to controls. (**D**) Sleep profile for young (top) and old (bottom, brown shading) (i) tim>2n4r tau^WT^, (ii) tim>0n4r tau^R460W^, (iii) Pdf>2n4r tau^WT^ and (iv) Pdf>0n4r tau^R406W^ flies compared to relevant controls. All genotypes show a normal bimodal pattern. (**E-F**) Quantifying daytime and night-time sleep, respectively. Tim>2n4r tau^WT^ expression does not affect daytime sleep relative to controls. Whereas tim>0n4r tau^R406W^ expression increases daytime sleep in young flies, but reduces daytime sleep in old flies, relative to controls. On the other hand, Pdf>tau expression does not affect daytime sleep (except for in young Pdf>2n4r tau^WT^ flies). But tim>tau and Pdf>tau expression greatly reduces night sleep compared to controls. Statistics described in Fig. 2 and Materials and Methods. n = 28-115 flies from 2-6 independent experiments.

We show tim-driven expression of tau^WT^ and tau^R406W^ produced a differential effect on daytime activity and sleep. As tim>2n4r tau^WT^ expression did not affect the level of activity or sleep during the day with respect to the controls in either young or old flies (Figs. 4B, Di, E), but tim>0n4r tau^R406W^ expression had an age-specific effect on daytime activity and sleep levels (Figs. 4B, Dii, E). Specifically, the young tim>0n4r tau^R406W^ flies exhibited reduced daytime activity and increased daytime sleep. Opposingly, the old tim>0n4r tau^R406W^ flies were more active and slept less during daytime than the controls (Figs 4B, E). In contrast, we found no statically significant age-related differences in daytime activity and sleep in either the tim>2n4r tau^WT^ or control flies (Figs. 4B, E). Hence, we have discovered isoform-specific differences on the clock-specific tau-mediated changes in total activity and sleep through the day, but not at night.

As the PDF clock-neurons are essential for controlling the timing of activity and sleep (Renn et al., 1999; Grima et al., 2004; Stoleru et al., 2004; Shang et al., 2008; Sheeba et al., 2008), we next asked whether PDF-positive pacemaker neuron restricted tau expression is sufficient to alter behavioural rhythms under LD conditions. The activity profiles revealed the Pdf>2n4r tau^WT^ (Fig. 4Av) and Pdf>0n4r tau^R406W^ (vi) flies exhibited normal bimodal rhythms with peaks around the light transitions but elevated basal activity. Hence unsurprisingly, their total night-activity was greatly increased (Fig. 4C) and night sleep-was severely reduced (Figs. 4Diii-iv, F) with respect to controls. However, the Pdf>tau flies’ daytime activity (Fig. 4B) and sleep levels (Figs. 4Diii-iv, E) were normal, except for a small daytime sleep gain in the young Pdf>2n4r tau^WT^ flies (Figs. 4Diii, E). These findings collectively show that the extent of tau expression has little effect on the night-activity gain and -sleep loss phenotype, as Elav>, tim> and Pdf> driven tau expression caused a similar night-activity increase and -sleep reduction. Hence, tau expression in the PDF clock-neurons promotes night activity and suppresses night sleep. And, tau expression in the non-PDF clock-neurons has neither an additive nor synergistic effect on the night-activity gain and -sleep loss phenotype.

### Tau expression in the PDF-positive pacemaker neurons fails to evoke DD behavioural arrhythmicity

Next, we tested whether pan-clock tau expression perturbs “free-running” circadian locomotor activity rhythms. Both the Gal4-(Fig. 5Ai) and UAS-control flies (i-ii) maintained a robust rhythm of daytime activity and night inactivity with a period of just under 24 h, comparable to wild-type flies (i) in DD conditions. However, in clear contrast, the tim>2n4r tau^WT^ (Fig. 5Aiii) and tim>0n4r tau^R406W^ (iv) flies failed to show any apparent potentiation between daytime activity and night inactivity, as relative night activity appeared to be substantially elevated. Instead, both young and old tim>tau flies exhibited similar highly significant reductions in DD behavioural rhythmicity with respect to their age-matched controls (Fig. 5B). A subpopulation of the old tim>2n4r tau^WT^ (∼30 %) and tim>0n4r tau^R406W^ (<5 %) flies developed arrhythmia, showing similar activity levels throughout the 24 h period. In comparison, all the old control flies remained rhythmic (Table. 1). Overall, DD rhythmicity declined as the flies aged; but the age-related dysrhythmia was similar in the tim>tau and control flies (Fig. 5B). We found that Elav>tau and tim>tau expression produced similar behavioural arrhythmicity, except for a larger arrhythmic tim>tau sub-population (Table. 1), suggesting that tau expression in the non-clock-neurons has neither an additive nor deleterious effect on the arrhythmic phenotype.

**Figure 5.**
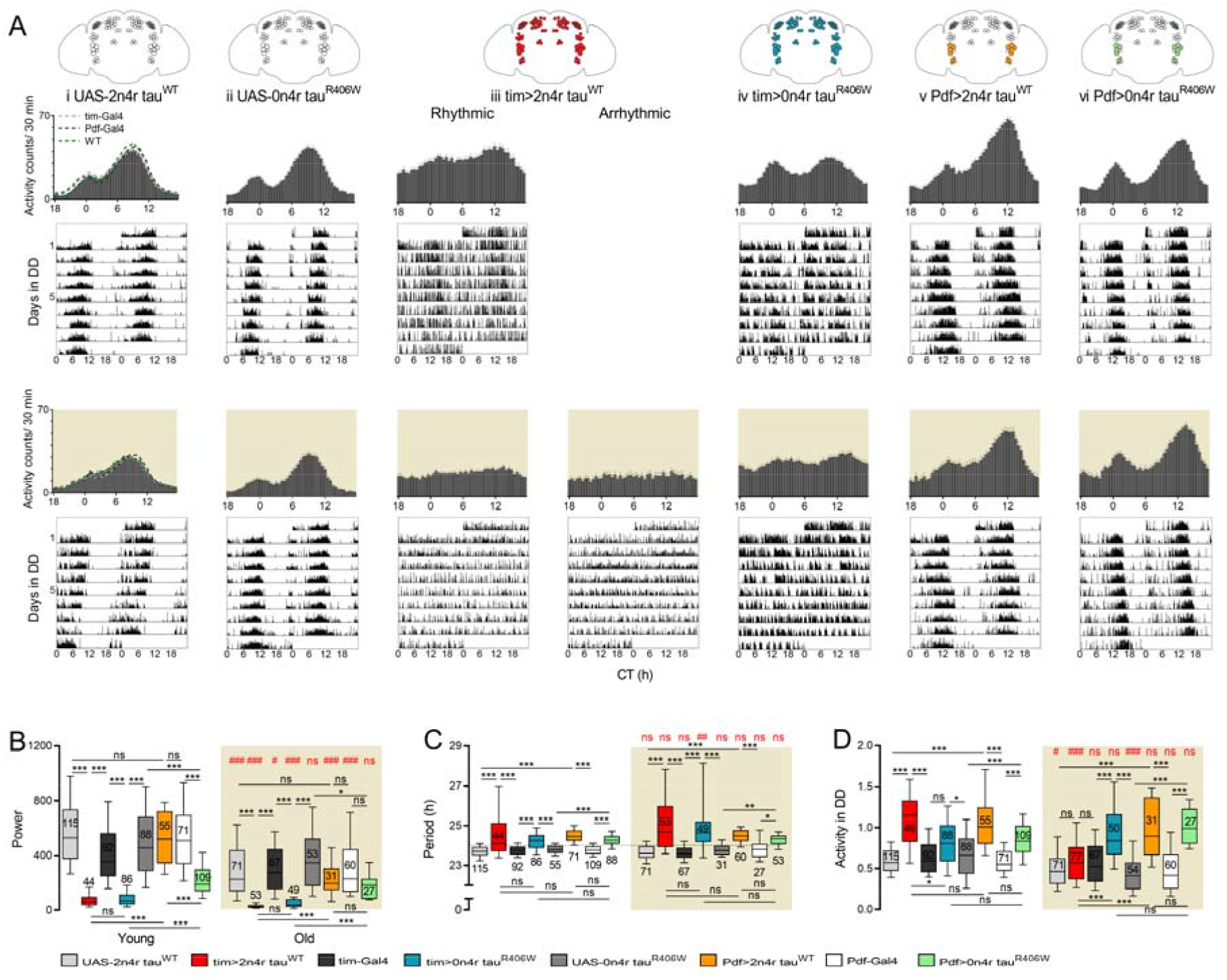
Pan-clock, but not central clock-neuron restricted, tau expression is sufficient to produce reduced DD rhythmicity. **(A)** Activity histograms and representative double-plotted actograms for 5-(young) (top) and 25-day old (old) (bottom, brown shading) (i-ii) UAS control, (iii) tim>2n4r tau^WT^, (iv) tim>0n4r tau^R406W^, (v) Pdf>2n4r tau^WT^ and (vi) Pdf>0n4r tau^R406W^ flies. Control flies maintain robust daytime activity and night-time inactivity rhythms, with a ∼24 h period. Tim>tau flies show weak rhythms with a long period (rightward shift in the activity peak) or are arrhythmic (for the arrhythmic fly % see Table. 1). Pdf>tau flies exhibit robust rhythms with a long period. (**B**) Greatly reduced DD rhythmicity in the tim>tau flies relative to control and Pdf>tau flies. (**C**) Prolongation of the behavioural period in DD in the tim>tau and Pdf>tau flies compared to controls. (**D**) Greatly increased overall activity in the tim>tau and Pdf>tau flies relative to controls (except for in young tim>0n4r tau^R406W^ and old tim>2n4r tau^WT^ flies). Statistics described in Fig. 3 and Materials and Methods. n = 27 – 115 flies from 2-6 independent experiments.

Interestingly, tim>tau expression, unlike Elav>tau expression (Figs. 3A, B), significantly increased the period length of DD rhythms (Fig. 5C and Table. 1), demonstrating a central clock defect. A rightward shift in the activity peak is visible in the histograms and actograms of the tim>tau flies (Figs. 5Ai-iv). Both tim>2n4r tau^WT^ and tim>0n4r tau^R406W^ expression produced similar prolongation of the behavioural period in DD. Furthermore, the tim>tau flies’ period lengths varied widely (Fig. 5C and Table. 1), an increase in variation is associated with both normal ageing and AD (Musiek et al., 2015). We found tim>tau expression also made the flies DD hyperactive, but an age-related loss of the hyperactive phenotype was seen in the tim>2n4r tau^WT^ flies (Fig. 5D). These results show that pan-clock tau expression produced very weak long-period rhythms with elevated activity in DD.

As the PDF clock-neurons are important for DD rhythmicity (Grima et al., 2004; Stoleru et al., 2004; Stoleru et al., 2005; Yao and Shafer, 2014), we asked whether PDF neuron tau expression was sufficient to disturb intrinsic circadian behaviour. Interestingly, Pdf>2n4r tau^WT^ expression exhibited robust circadian behaviour, where DD rhythmicity did not significantly differ from the young and old controls (Figs. 5A, B). Moreover, Pdf>0n4r tau^R406W^ flies maintained obvious behavioural rhythmicity. Visible inspection of the histograms and actograms revealed most individuals upheld a discernible pattern of 12 h of activity followed by 12 h of inactivity, similar to control flies (Fig. 5Avi). However, the Pdf>0n4r tau^R406W^ flies’ behavioural rhythmicity was reduced, compared to the controls, although the difference failed to reach the significance threshold in old flies (Fig. 5B). These results indicate that PDF clock-neuron tau expression is insufficient to affect circadian rhythmicity in DD and identify the non-PDF clock-neurons as the major drivers of tau-related DD behavioural arrhythmia.

However, we found that the Pdf>tau flies exhibited behavioural changes despite the normal behavioural rhythmicity. Specifically, Pdf>tau expression generated long-period DD rhythms. We observed no statistically significant age-, region- or isoform-specific differences on the period-lengthening effects (Figs. 5A, C and Table. 1). We found that Pdf>tau expression also produced an overall increase in DD activity, and the hyperactive phenotype was not significantly affected by either isoform or age (Fig. 5D). These results show that PDF clock-neuron tau expression is sufficient to prolong the behavioural period and induce hyperactivity. Varying the extent of clock-restricted tau^WT^, but not tau^R406W^, expression significantly affected the DD hyperactive phenotype, specifically in old flies. This change resulted from the age-related loss of the elevated overall activity in the tim>2n4r tau^WT^ flies (Figs. 5A, D). These results show that the PDF clock-neurons are the main site of the tau-mediated period-lengthening and hyperactive phenotypes.

### Behavioural arrhythmicity is not due to disruption of the molecular clock

Next, we investigated whether the circadian behavioural arrhythmia in the pan-clock tau-expressing flies coincides with damage to the molecular clock. To answer this question, we monitored period oscillations in the clock-neurons by recording bioluminescence from a per luciferase fusion construct (Stanewsky et al., 1997). Comparing tim>2n4r tau^WT^ and control flies, we found similar bioluminescence oscillations (Fig. 6A). Rhythmicity, as assessed by the relative amplitude error (RAE) that varies from 0 (strong rhythm) to 1 (arrhythmic/no rhythm), was similar among the tim>tau and control flies (Fig. 6B). No statistically significant age-related decline was seen in the strength of molecular rhythms (Fig. 6B), unlike in behavioural rhythms (Figs. 5A, B), in either the tim>tau or control flies. The period length of the oscillations in the tim>tau flies did not differ significantly from age-matched controls, despite increased variation in the tim>tau flies (Fig. 6C). These results show that the molecular clock remains functional in the tau-expressing flies and suggests that perturbed clock-neuron output or communication drives the behavioural arrhythmicity.

**Figure 6.**
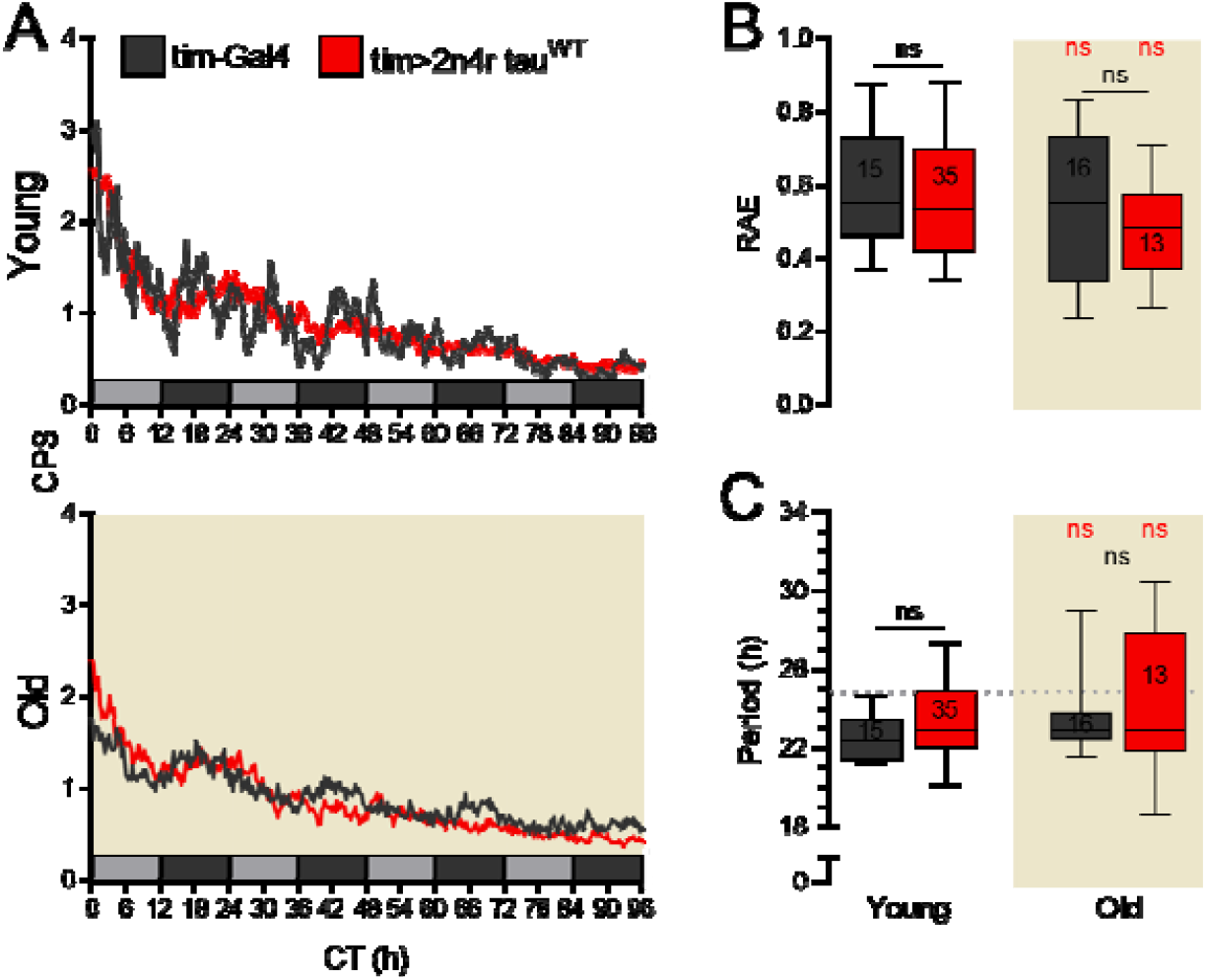
Intact molecular clock despite the behavioural arrhythmia in the tim>tau flies. (**A**) Bioluminescence profiles for 5-(young) (top) and 25-day old (old) (bottom, brown shading) control and tim>2n4r tau^W^^T^ flies during 4 days of DD. Robust Per oscillations are seen in control and tim>tau flies. (**B**-**C**) No statistically significant differences in RAE or period are found between tim>tau and control flies in both age groups. Comparison between different genotypes of the same age (black asterisks) and different ages of the same genotype (red number symbols) by Mann-Whitney U-tests. n= 13-35 flies from 2 independent experiments.

### Tau-related behavioural arrhythmia is independent of the PDF-positive pacemaker neurons

We have shown both broad (Fig. 3) and PDF clock-neuron restricted (Fig. 5) tau expression results in flies that exhibit similar night-activity gains and -sleep loss. To determine whether these behaviour changes are attributable to tau expression within the central clock-neurons, we used a Pdf-Gal80 transgene, which blocks Gal4 activity in the PDF clock-neurons (Stoleru et al., 2004). The activity profiles of the young Elav, Pdf-Gal80>0n4r tau^R406W^ and control flies were strikingly similar, with greater activity during the day than at night (Fig. 7A). Unsurprisingly, the daytime and night-time activity and sleep levels did not significantly differ between the Elav, Pdf-Gal80>0n4r tau^R406W^ and control flies (Figs. 7B-C). Therefore, blocking tau expression only in the PDF clock-neurons fully rescues the tau-related night-activity gain and -sleep loss phenotype, indicating these behavioural changes originate from tau within the central clock-neurons.

**Figure 7.**
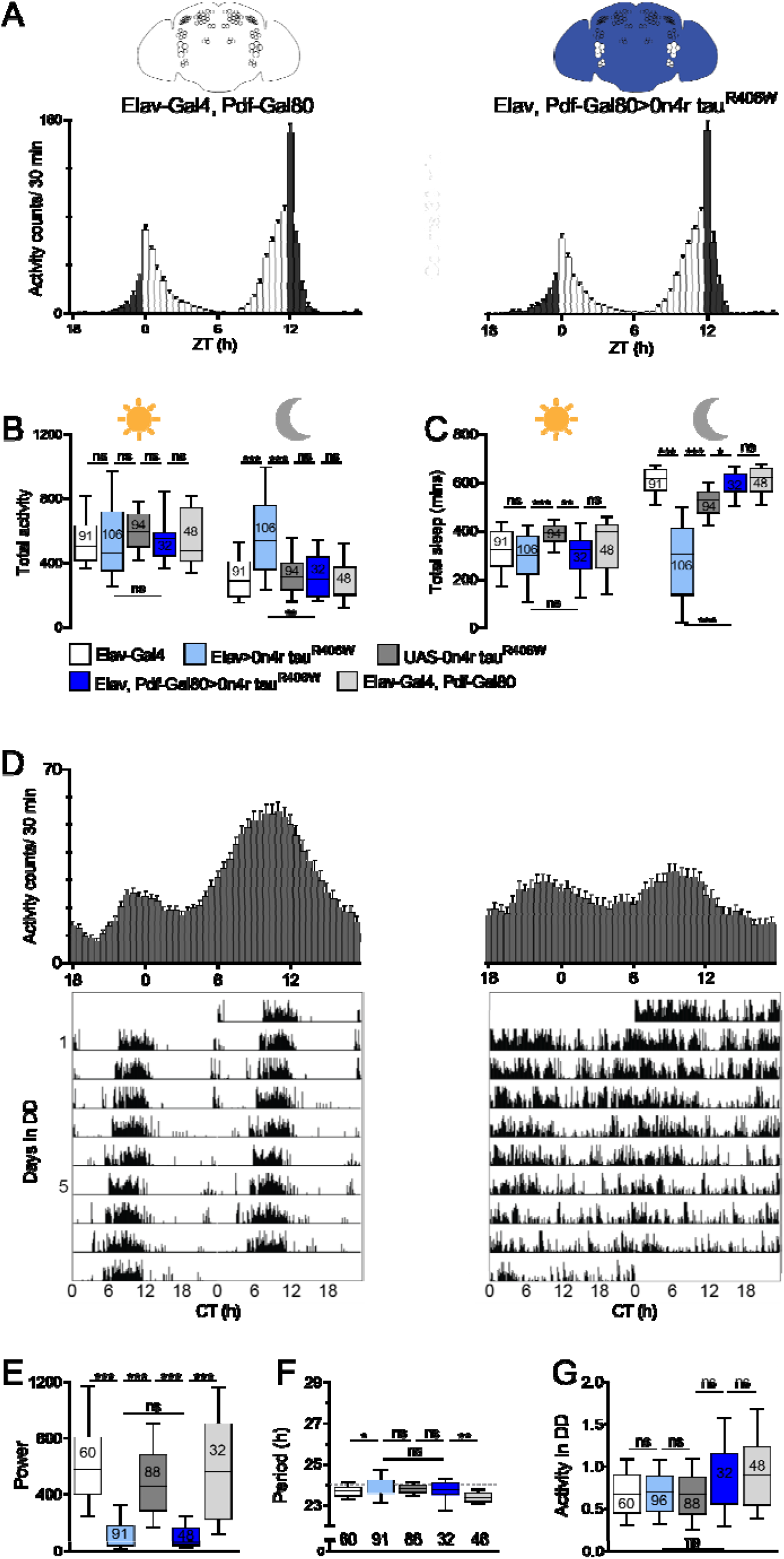
Blocking tau expression in the central clock rescues the altered activity and sleep levels under LD conditions, but not the reduced rhythmicity in DD conditions. (**A**) Activity histograms for 5-day old (young) control and Elav, Pdf-Gal80> 0n4r tau^R406W^ flies under LD conditions. Both control and Elav, Pdf-Gal80> 0n4r tau^R406W^ flies show normal bimodal activity rhythms. (**B**) No change in daytime and night activity in Elav, Pdf-Gal80>0n4r tau^R406W^ flies relative to controls. (**C**) No change in daytime and night sleep in Elav, Pdf-Gal80>0n4r tau^R406W^ flies relative to controls. (**D**) Average activity histograms and representative actograms for young control and Elav, Pdf-Gal80>0n4r tau^R406W^ flies. Elav, Pdf-Gal80>0n4r tau^R406W^ flies show very weak rhythms with a normal period. (**E**) Greatly reduced DD rhythmicity in the Elav, Pdf-Gal80>0n4r tau^R406W^ flies relative to controls. (**F**) Elav, Pdf-Gal80>0n4r tau^R406W^ expression did not alter the behavioural period in DD. (**G**) Elav, Pdf, Gal80>0n4r tau^R406W^ and control flies show comparable overall activity levels. (**B, E, G**) Multiple comparisons between different genotypes by 1-way ANOVA and post-hoc Tukey HSD tests with log-transformed data. (**C, F**) Multiple comparisons between different genotypes by Kruskal-Wallis ANOVA and post-hoc Dunn’s tests. (**B-C**) 32–106 flies from 2-6 independent experiments. (**E-G**) 32-91 flies from 2-6 independent experiments.

As Pdf>tau expression failed to disrupt DD rhythmic behaviour (Fig. 5), we next investigated whether restricting PDF clock-neuron tau expression was sufficient to ameliorate the arrhythmic phenotype. In the young Elav Pdf-Gal80>0n4r tau^R406W^ flies, activity during the subjective night appeared to be elevated, resulting in a reduced day/night difference in activity (Fig. 7D). Hence, unsurprisingly their DD behavioural rhythmicity was severely reduced with respect to the controls (Fig. 7E). Overall, Elav, Pdf-Gal80> and Elav> driven tau expression produced almost identical changes in circadian behaviour; the rhythmicity (Fig. 7E), period (Fig. 7F) and 24 h activity (Fig. 7G) of locomotor behaviour in the Elav, Pdf-Gal80>0n4r tau^R406W^ flies did not differ statistically from the Elav>0n4r tau^R406W^ flies. Together, these results demonstrate that restricting PDF clock-neuron tau expression is insufficient to rescue the tau-related dysrhythmia. Hence, tau expression in the central clock-neurons is not necessary to cause behavioural arrhythmia, further highlighting that dysfunction in neuronal populations afferent to the PDF-positive pacemaker neurons is the primary driver of the tau-mediated arrhythmic phenotype.

## Discussion

### Pan-neuronal tau expression disrupts circadian behaviour under LD and DD conditions

Whilst circadian dysfunction is widespread in tauopathies, including AD (Satlin et al., 1995; Harper et al., 2001; Volicer et al., 2001), PD (Mantovani et al., 2018) and FTD (Harper et al., 2001; Anderson et al., 2009), it has been an open question whether tau misexpression can give rise to circadian behaviour deficits. To systematically assess the consequences of human tau expression in specific neuronal populations on behavioural and molecular rhythms, we first examined the effect on circadian behaviour of pan-neurally expressing human tau in the *Drosophila* brain. We found that broad neuronal human tau expression affected circadian locomotor activity rhythms under LD and DD conditions in young (∼5-day-old) and old (∼25-day-old) flies.

Firstly, under LD conditions, the Elav>tau flies showed bimodal activity rhythms but exhibited elevated activity during the night and a reduced day/night difference in activity (Figs. 2A-C). Tg4510 mice, which express the FTD-associated tau mutant, P301L, driven by the forebrain-specific CaMKIIα promoter, were more active during the day (*i*.*e*. the inactive phase) than control littermates (Stevanovic et al., 2017). Thus, both models reproduce the shift towards a higher proportion of the total activity occurring during the inactive phase, often seen in human AD patients (Volicer et al., 2001; Harper et al., 2004). Overall, our Elav>tau flies also slept less throughout the day and night (daytime sleep loss was statistically insignificant) (Figs. 2D-F). Adulthood-restricted pan-neuronal expression of Aβ^42^ yielded flies, which exhibited reduced and fragmented night sleep (Tabuchi et al., 2015). Similarly, young (2-3-day-old) Elav>Aβ^42^ flies displayed reduced total sleep (Gerstner et al., 2017). As such, broad neuronal tau and Aβ^42^ expression both produced a night-sleep loss. Hence, in AD, the presence of both tau and amyloid pathology may additively or synergistically disrupt sleep. In alignment with these data, AD patients’ sleep has often been reported to be reduced and fragmented at night (Prinz et al., 1982; Vitiello et al., 1990). Notably, neither Elav>tau (Figs. 2D-E) nor Elav>Aβ^42^ flies (Tabuchi et al., 2015; Gerstner et al., 2017) recapitulated the increased daytime drowsiness often reported in patients with AD (Volicer et al., 2001; Bliwise, 2004; Anderson et al., 2009).

Secondly, in “free-running” DD continuous darkness, pan-neuronal expression of tau generated progressive behavioural arrhythmia, as evidenced by reduced overall rhythmicity (Figs. 3A-B) and an increased arrhythmic sub-population (Table. 1), indicating an intrinsic circadian rhythm defect. These circadian rhythm perturbations are already present in young flies (∼5-day old). Therefore, they likely precede the onset of neurodegeneration; first identified in ∼10-day old Elav>0n4r tau^R406W^ flies (Wittmann et al., 2001). The dampening of circadian locomotor activity rhythms seen in our model is similar to that reported in AD (Volicer et al., 2001), FTD (Harper et al., 2001; Anderson et al., 2009) and PD (van Hilten et al., 1993; Placidi et al., 2008) patients.

### Clock-specific tau expression alters activity and sleep under LD conditions

Next, we investigated how clock-restricted tau expression affected circadian behaviour. We discovered that tau expression in either all the clock cells or the PDF clock-neurons similarly made the flies more active and sleep less during the night (Fig. 4). Additionally, blocking PDF neuron tau expression was sufficient to rescue the behavioural changes fully, as tau expression in all neurons except the PDF clock-neurons failed to alter total activity or sleep levels or the day/night difference in activity (Figs. 7A-C). As such, tau expression within the PDF clock-neurons was sufficient, and necessary, to produce the night-activity gain and -sleep loss phenotype. Hence, these behaviour changes arise from tau expression within the PDF clock-neurons, rather than the non-PDF clock-neurons or non-clock-neurons. The elevated night activity in the Pdf>tau flies is not attributable to a loss of PDF signalling, as Pdf null mutants have an advanced evening activity peak, which increases daytime activity (Renn et al., (1999).

### Targeted tau expression in the clock-network differentially affects circadian rhythms in DD conditions

We discovered restricting tau expression to the PDF clock-neurons, but not all clock-neurons, rescues the reduced DD behavioural rhythmicity (Fig. 5). Tau expression in all neurons except the PDF neurons yielded flies, which exhibited similar reductions in behavioural rhythmicity to the Elav>tau flies (Figs. 7D-E). As such, tau expression in the PDF clock-neurons is neither necessary nor sufficient to produce reduced DD rhythmicity. Hence, we have identified the non-PDF clock-neurons as the main site of the tau-related arrhythmic phenotype. One possible explanation is that tau disrupts the non-PDF clock-neurons’ (*i*.*e*. DN1 neurons) communication with non-clock output neurons (*i*.*e*. DH44 positive PI cells), which are necessary for behavioural rhythms (Cavanaugh et al., 2014; King et al., 2017)). In the tim>tau flies, we observed an intact molecular clock; as Per oscillations were not disturbed (Fig. 6), despite the behavioural arrhythmia (Figs. 5A-B). Similarly, the primary target for behavioural arrhythmia in Aβ^42^-expressing *Drosophila* (Chen et al., 2014; Long et al., 2014), R6/2 HD mice (Pallier et al., 2007) and AD patients (Wu and Swaab, 2007; Cermakian et al., 2011) is reported to be downstream of the central clock.

Clock tau expression yielded flies, which exhibited prolongation of the behaviour period in DD (Figs. 5A, C). These long-period rhythms cannot result from abolished PDF signalling, because chemical or genetic ablation of the PDF clock-neurons results in short-period rhythms (Renn et al. (1999). Furthermore, the long-period rhythms cannot be related to blocked chemical neurotransmission as Pdf>TNT flies show a similar behavioural period in DD to control flies (Umezaki et al., 2011). In DD, Tg4510 mice, which express a high level of tau^P301L^ in the entire forebrain, exhibited a ∼1 h longer period than control littermates (Stevanovic et al., 2017). Whereas we found clock-specific, but not broad neuronal, tau expression produced prolongation of the behavioural period. Hence, we showed that tau expression within the non-clock-neurons suppresses the period-lengthening effect of PDF clock-neurons tau expression. These different findings could be attributable to the different model organisms’ peculiarities or the differential effects of tau^P301L^ and tau^WT^/tau^R406W^.

We found that Pdf> and tim>tau expression similarly increased overall DD activity (Fig. 5D). Hence, tau’s activity-promoting effect arises from within the PDF clock-neurons, rather than the non-PDF clock-neurons. Because Pdf>tau (Fig. 5D), but not Elav>tau (Fig. 3D), expression produced DD hyperactivity, tau expression in the non-clock-neurons attenuates the activity-promoting effect of PDF clock-neuron tau expression. For example, tau expression might interfere with communication between the central clock and brain regions involved in locomotion control (*e*.*g*. the ellipsoid body). As a result of wandering, most human AD patients exhibit elevated activity (Logsdon et al., 1998). Other studies have shown electrophysiological PDF clock-neuron abnormalities coincide with gains in activity and loss of sleep in disease models (Sheeba et al., 2008). Accordingly, in our model, specific behaviour changes likely arise from neurophysiological changes in either the PDF or non-PDF clock-neurons.

### Human tau expression in *Drosophila* faithfully recapitulates the human AD sleep and circadian rhythm defects

Here we showed that tau expression in the *Drosophila* brain causes circadian abnormalities closely matching those found in human AD and other tauopathy patients. These results validate the use of *Drosophila* as a model to study the effects of tau pathology on circadian behaviour. We described the clock neuronal subpopulations that mediate discrete circadian behavioural deficits and specifically identified the PDF clock-neurons as the main site of behavioural changes in the overall amount of activity and sleep (restricted to the LD night) (Figs 4, 7A-C). We further identified the non-PDF clock-neurons as the main site of activity distribution changes (Figs. 5, 7D-G) and discovered that the circadian behavioural deficits arise from clock-neuron dysfunction, rather than death; as shown by ongoing Per oscillations (Fig. 6). The fly model we described in this study provides the opportunity to study the circuitry that mediates tau-related circadian and sleep deficits. Further understanding will hopefully enable the development of novel therapeutics that improve well-being and clinical outcome in patients.

## Acknowledgements

This work was supported by Biotechnology and Biological Sciences Research Council BB/H013849/1, BB/F012071/1 and BB/D001900/1, Engineering and Physical Sciences Research Council EP/P006094/1, Leverhulme Trust RPG-2012–567, Jane and Aatos Erkko Foundation, High-End Foreign Expert Grant by Chinese Government GDT20051100004 and Beijing Normal University Open Research Fund to M.J.; Biotechnology and Biological Sciences Research Council BB/M000435/1 and BB/N018540/1 and 111 Project # D16014 to S.J.D.; We thank Mel Feany, Ralf Stanewsky and Charlotte Forster for *Drosophila*; Amanda M. Davis for luminescence test support; and Cahir O’Kane and members of the M.J. laboratory for discussions.

## References

Ali YO, Ruan K, Zhai RG (2012) NMNAT suppresses Tau-induced neurodegeneration by promoting clearance of hyperphosphorylated Tau oligomers in a Drosophila model of tauopathy. Hum Mol Genet 21:237–250.

Alzheimer A (1906) Über einen eigenartigen schweren Erkrankungsprozeß der Hirnrinde.. Neurologisches Centralblatt 23:1129–1136.

Anderson KN, Hatfield C, Kipps C, Hastings M, Hodges JR (2009) Disrupted sleep and circadian patterns in frontotemporal dementia. Eur J Neurol 16:317–323.

Anwer MU, Boikoglou E, Herrero E, Hallstein M, Davis AM, James GV, Nagy F, Davis SJ (2014) Natural variation reveals that intracellular distribution of ELF3 protein is associated with function in the circadian clock. Elife 3.

Bell-Pedersen D, Cassone VM, Earnest DJ, Golden SS, Hardin PE, Thomas TL, Zoran MJ (2005) Circadian rhythms from multiple oscillators: Lessons from diverse organisms. Nat Rev Genet 6:544–556.

Bliwise DL (2004) Sleep disorders in Alzheimer’s disease and other dementias. In: Clinical Cornerstone, pp 16–28.

Braak H, Braak E (1988) Neuropil Threads Occur in Dendrites of Tangle-Bearing Nerve-Cells. Neuropath Appl Neuro 14:39–44.

Brand AH, Perrimon N (1993) Targeted Gene-Expression as a Means of Altering Cell Fates and Generating Dominant Phenotypes. Development 118:401–415.

Buhr E, Takahashi JS (2013) Genetic control of the circadian pacemaker. Genetic Basis of Sleep and Sleep Disorders:119–126.

Cavanaugh DJ, Geratowski JD, Wooltorton JRA, Spaethling JM, Hector CE, Zheng XZ, Johnson EC, Eberwine JH, Sehgal A (2014) Identification of a Circadian Output Circuit for Rest: Activity Rhythms in Drosophila. Cell 157:689–701.

Cermakian N, Lamont EW, Boudreau P, Boivin DB (2011) Circadian Clock Gene Expression in Brain Regions of Alzheimer’s Disease Patients and Control Subjects. J Biol Rhythm 26:160–170.

Chen KF, Possidente B, Lomas DA, Crowther DC (2014) The central molecular clock is robust in the face of behavioural arrhythmia in a Drosophila model of Alzheimer’s disease. Dis Model Mech 7:445–458.

Donlea JM, Pimentel D, Miesenbock G (2014) Neuronal Machinery of Sleep Homeostasis in Drosophila. Neuron 81:860–872.

Dubowy C, Sehgal A (2017) Circadian Rhythms and Sleep in Drosophila melanogaster. Genetics 205:1373–1397.

Gerstner JR, Lenz O, Vanderheyden WM, Chan MT, Pfeiffenberger C, Pack AI (2017) Amyloid-induces sleep fragmentation that is rescued by fatty acid binding proteins in Drosophila. J Neurosci Res 95:1548–1564.

Grima B, Chelot E, Xia RH, Rouyer F (2004) Morning and evening peaks of activity rely on different clock neurons of the Drosophila brain. Nature 431:869–873.

Harper DG, Stopa EG, McKee AC, Satlin A, Fish D, Volicer L (2004) Dementia severity and Lewy bodies affect circadian rhythms in Alzheimer disease. Neurobiol Aging 25:771–781.

Harper DG, Stopa EG, McKee AC, Satlin A, Harlan PC, Goldstein R, Volicer L (2001) Differential circadian rhythm disturbances in men with Alzheimer disease and frontotemporal degeneration. Arch Gen Psychiat 58:353–360.

Hendricks JC, Finn SM, Panckeri KA, Chavkin J, Williams JA, Sehgal A, Pack AI (2000) Rest in Drosophila is a sleep-like state. Neuron 25:129–138.

Hutton M et al. (1998) Association of missense and 5 ‘-splice-site mutations in tau with the inherited dementia FTDP-17. Nature 393:702–705.

Jackson GR, Wiedau-Pazos M, Sang TK, Wagle N, Brown CA, Massachi S, Geschwind DH (2002) Human wild-type tau interacts with wingless pathway components and produces neurofibrillary pathology in Drosophila. Neuron 34:509–519.

Josephs KA (2017) Current Understanding of Neurodegenerative Diseases Associated With the Protein Tau. Mayo Clin Proc 92:1291–1303.

Kaneko M, Hall JC (2000) Neuroanatomy of cells expressing clock genes in Drosophila: Transgenic manipulation of the period and timeless genes to mark the perikarya of circadian pacemaker neurons and their projections. J Comp Neurol 422:66–94.

Kauranen H, Menegazzi P, Costa R, Helfrich-Forster C, Kankainen A, Hoikkala A (2012) Flies in the North: Locomotor Behavior and Clock Neuron Organization of Drosophila montana. J Biol Rhythm 27:377–387.

King AN, Barber AF, Smith AE, Dreyer AP, Sitaraman D, Nitabach MN, Cavanaugh DJ, Sehgal A (2017) A Peptidergic Circuit Links the Circadian Clock to Locomotor Activity. Curr Biol 27:1915-+.

Koss DJ, Robinson L, Drever BD, Plucinska K, Stoppelkamp S, Veselcic P, Riedel G, Platt B (2016) Mutant Tau knock-in mice display frontotemporal dementia relevant behaviour and histopathology. Neurobiol Dis 91:105–123.

Locke JCW, Southern MM, Kozma-Bognar L, Hibberd V, Brown PE, Turner MS, Millar AJ (2005) Extension of a genetic network model by iterative experimentation and mathematical analysis. Mol Syst Biol 1.

Logsdon RG, Teri L, McCurry SM, Gibbons LE, Kukull WA, Larson EB (1998) Wandering: A significant problem among community-residing individuals with Alzheimer’s disease. J Gerontol B-Psychol 53:P294–P299.

Long DM, Blake MR, Dutta S, Holbrook SD, Kotwica-Rolinska J, Kretzschmar D, Giebultowicz JM (2014) Relationships between the Circadian System and Alzheimer’s Disease-Like Symptoms in Drosophila. Plos One 9.

Mantovani S, Smith SS, Gordon R, O’Sullivan JD (2018) An overview of sleep and circadian dysfunction in Parkinson’s disease. J Sleep Res 27.

Means JC, Venkatesan A, Gerdes B, Fan JY, Bjes ES, Price JL (2015) Drosophila Spaghetti and Doubletime Link the Circadian Clock and Light to Caspases, Apoptosis and Tauopathy. Plos Genet 11.

Mershin A, Pavlopoulos E, Fitch O, Braden BC, Nanopoulos DV, Skoulakis EMC (2004) Learning and memory deficits upon TAU accumulation in Drosophila mushroom body neurons. Learn Memory 11:277–287.

Musiek ES, Xiong DD, Holtzman DM (2015) Sleep, circadian rhythms, and the pathogenesis of Alzheimer Disease. Exp Mol Med 47.

Nishimura I, Yang YF, Lu BW (2004) PAR-1 kinase plays an initiator role in a temporally ordered phosphorylation process that confers tau toxicity in Drosophila. Cell 116:671–682.

Pallier PN, Maywood ES, Zheng ZG, Chesham JE, Inyushkin AN, Dyball R, Hastings MH, Morton AJ (2007) Pharmacological imposition of sleep slows cognitive decline and reverses dysregulation of circadian gene expression in a transgenic mouse model of huntington’s disease. J Neurosci 27:7869–7878.

Placidi F, Izzi F, Romigi A, Stanzione P, Marciani MG, Brusa L, Sperli F, Galati S, Pasqualetti P, Pierantozzi M (2008) Sleep-wake cycle and effects of cabergoline monotherapy in de novo Parkinson’s disease patients - An ambulatory polysomnographic study. J Neurol 255:1032–1037.

Prinz PN, Vitaliano PP, Vitiello MV, Bokan J, Raskind M, Peskind E, Gerber C (1982) Sleep, Eeg and Mental Function Changes in Senile Dementia of the Alzheimer Type. Neurobiol Aging 3:361–370.

Renn SCP, Park JH, Rosbash M, Hall JC, Taghert PH (1999) A pdf neuropeptide gene mutation and ablation of PDF neurons each cause severe abnormalities of behavioral circadian rhythms in Drosophila. Cell 99:791–802.

Saito Y, Geyer A, Sasaki R, Kuzuhara S, Nanba E, Miyasaka T, Suzuki K, Murayama S (2002) Early-onset, rapidly progressive familial tauopathy with R406W mutation. Neurology 58:811–813.

Salmon DP, Heindel WC, Lange KL (1999) Differential decline in word generation from phonemic and semantic categories during the course of Alzheimer’s disease: Implications for the integrity of semantic memory. J Int Neuropsych Soc 5:692–703.

Satlin A, Volicer L, Stopa EG, Harper D (1995) Circadian Locomotor-Activity and Core-Body Temperature Rhythms in Alzheimers-Disease. Neurobiol Aging 16:765–771.

Shang YH, Griffith LC, Rosbash M (2008) Light-arousal and circadian photoreception circuits intersect at the large PDF cells of the Drosophila brain. P Natl Acad Sci USA 105:19587–19594.

Shaw PJ, Cirelli C, Greenspan RJ, Tononi G (2000) Correlates of sleep and waking in Drosophila melanogaster. Science 287:1834–1837.

Sheeba V, Fogle KJ, Kaneko M, Rashid S, Chou YT, Sharma VK, Holmes TC (2008) Large Ventral Lateral Neurons Modulate Arousal and Sleep in Drosophila. Curr Biol 18:1537–1545.

Stanewsky R, Jamison CF, Plautz JD, Kay SA, Hall JC (1997) Multiple circadian-regulated elements contribute to cycling period gene expression in Drosophila. Embo J 16:5006–5018.

Stevanovic K, Yunus A, Joly-Amado A, Gordon M, Morgan D, Gulick D, Gamsby J (2017) Disruption of normal circadian clock function in a mouse model of tauopathy. Exp Neurol 294:58–67.

Stoleru D, Peng Y, Agosto J, Rosbash M (2004) Coupled oscillators control morning and evening locomotor behaviour of Drosophila. Nature 431:862–868.

Stoleru D, Peng Y, Nawathean P, Rosbash M (2005) A resetting signal between Drosophila pacemakers synchronizes morning and evening activity. Nature 438:238–242.

Swaab DF, Fliers E, Partiman TS (1985) The Suprachiasmatic Nucleus of the Human-Brain in Relation to Sex, Age and Senile Dementia. Brain Res 342:37–44.

Swaab DF, Lucassen PJ, Salehi A, Scherder EJA, van Someren EJW, Verwer ARWH (1998) Reduced neuronal activity and reactivation in Alzheimer’s disease. Prog Brain Res 117:343–377.

Tabuchi M, Lone SR, Liu S, Liu QL, Zhang JL, Spira AP, Wu MN (2015) Sleep Interacts with A beta to Modulate Intrinsic Neuronal Excitability. Curr Biol 25:702–712.

Umezaki Y, Yasuyama K, Nakagoshi H, Tomioka K (2011) Blocking synaptic transmission with tetanus toxin light chain reveals modes of neurotransmission in the PDF-positive circadian clock neurons of Drosophila melanogaster. J Insect Physiol 57:1290–1299.

van Dongen AMJ, Chapman ML (1999) K channel gating by a dynamic selectivity filter. Biophys J 76:A415–A415.

van Hilten JJ, Weggeman M, Vandervelde EA, Kerkhof GA, Vandijk JG, Roos RAC (1993) Sleep, Excessive Daytime Sleepiness and Fatigue in Parkinsons-Disease. J Neural Transm-Park 5:235–244.

Vitiello MV, Prinz PN, Williams DE, Frommlet MS, Ries RK (1990) Sleep disturbances in patients with mild-stage Alzheimer’s disease. J Gerontol 45:M131–138.

Volicer L, Harper DG, Manning BC, Goldstein R, Satlin A (2001) Sundowning and circadian rhythms in Alzheimer’s disease. Am J Psychiat 158:704–711.

Wittmann CW, Wszolek MF, Shulman JM, Salvaterra PM, Lewis J, Hutton M, Feany MB (2001) Tauopathy in Drosophila: Neurodegeneration without neurofibrillary tangles. Science 293:711–714.

Wu YH, Swaab DF (2007) Disturbance and strategies for reactivation of the circadian rhythm system in aging and Alzheimer’s disease. Sleep Med 8:623–636.

Yao Z, Shafer OT (2014) The Drosophila Circadian Clock Is a Variably Coupled Network of Multiple Peptidergic Units. Science 343:1516–1520.

Zhou JN, Hofman MA, Swaab DF (1995) Vip Neurons in the Human Scn in Relation to Sex, Age, and Alzheimers-Disease. Neurobiol Aging 16:571–576.

